# Divergent Side-Chain Networks of the Mineralocorticoid Receptor Control Mutation-Altered Drug Response

**DOI:** 10.64898/2025.12.18.695081

**Authors:** Johannes Jokiel, Marcel Bermúdez

## Abstract

The mineralocorticoid receptor (MR) is a nuclear hormone receptor whose activation depends on ligand-induced conformational changes of the activation function-2, particularly involving helix 12. The rare gain-of-function mutation S810L was reported to alter MR pharmacology by converting classical steroidal antagonists such as spironolactone or eplerenone into agonists. In contrast, the non-steroidal antagonist finerenone has been shown to retain its functionality. However, the mechanistic reason for those ligand-specific effects remains elusive. To uncover the molecular basis for this ligand-specific behavior, we performed molecular dynamics simulations of wild-type and S810L MR bound to eplerenone or finerenone. We combined distance metrics with a residue-centered dynophore analysis and MDPath to track allosteric communication pathways and time-resolved side-chain interactions. In S810L, eplerenone shifts AF-2 side-chain networks toward H12, consistent with an agonist-like topology. Finerenone, by contrast, induces a local H3-H5 compaction around M777-810 yet diverts the AF-2 network away from H12, thereby preserving an antagonist-like topology. These network reallocations are absent in wild-type simulations, indicating a mutation-specific ligand-depending divergence. Together, these findings provide a mechanistic explanation for ligand-specific antagonist resilience in S810L MR and represent an example for structural pharmacogenomic approaches.

## 1 Introduction

The mineralocorticoid receptor (MR) is an important cardiovascular drug target, and mineralocorticoid receptor antagonists such as spironolactone, eplerenone and, more recently, finerenone are widely used to treat hypertension, heart failure and chronic kidney disease (Pitt et al. 1999; Kolkhof et al. 2014). Among the rare genetic variants of MR, there is a specific gain-of-function mutation, S810L, that causes a severe early-onset familial hypertension which worsens during pregnancy (Geller et al. 2000). Functional studies have shown that S810L shifts MR into an active-like state with elevated basal activity in the absence of aldosterone and induces a paradoxical shift in the response to steroidal MR antagonists such as spironolactone and eplerenone, which no longer behave as pure antagonists but exhibit partial agonist activity and further enhance MR signaling (Hultman et al. 2005).

Like other nuclear receptors, MR comprises a conserved ligand-binding domain (LBD) that undergoes hormone-dependent conformational changes. Agonist binding stabilizes the LBD in a conformation where helix 12 (H12) is positioned over the ligand pocket. Recent in-silico studies further detailed these ligand-dependent conformational rearrangements within the MR-LBD (Liang et al. 2025). This rearrangement generates the activation function-2 (AF-2) surface required for coactivator recruitment and transcriptional activation (Bourguet et al. 2000). In this state, helices 3-5 and 12 form a hydrophobic cleft that binds coactivator proteins. Antagonists alter this process through distinct mechanisms. Spironolactone and eplerenone promote a partial agonist-like state in which H12 adopts a closed position similar to the agonistic H12 position but in a destabilized, more inactive conformation (Hultman et al. 2005). This so-called *passive* antagonism can further be promoted by rapid ligand dissociation, mobilizing H12 (Amazit et al. 2015). By co-expressing the MR-LBD with a C-terminal, thrombin-cleavable NCOA1 peptide, and removing the covalent linker before crystallization, the elperenone-bound MR was structurally determined with closed H12 conformation (Bamberg et al. 2018). By contrast, *active* antagonists often contain bulky substituents that protrude toward H12 and sterically displace H12 from the coactivator site (Amazit et al. 2015). In both modes, antagonists interfere with AF-2 formation highlighting the central role of H12 positioning in MR regulation.

Classic MR antagonists like spironolactone and eplerenone are steroidal spirolactone derivatives and are widely prescribed for the treatment of hypertension, heart failure and other cardiovascular conditions. Spironolactone, a first-generation MR blocker also interacts with other steroid hormone receptors, which results in endocrine side effects such as gynecomastia and menstrual irregularities. Eplerenone is a second-generation antagonist that offers improved receptor selectivity, but still presents similar side effects and moderate potency. Finerenone is a novel dihydropyridine-based MR antagonist that combines high affinity and selectivity with reduced side effects. Finerenone showed greater organ-protective efficacy and less hyperkalemia than steroidal antagonists (Kolkhof et al. 2014). Structurally, it differs from spirolactone derivatives: when bound to MR it extends its polar carboxamide and bulky rings deeper toward H12. This orientation further allows dual formation of specific contacts with key residues A773 (H3) and S810 (H5), a mode of engagement not reported for steroid ligands. Mutating these residues with their counterparts from the glucocorticoid and progesterone receptors (A773G and S810M), drastically reduce finerenone’s potency. These insights emphasize the essential contribution of these side chains to ligand recognition and affinity of finerenone (Amazit et al. 2015).

The S810L mutation is located at the interface of H5 and H3 in the LBD and represents a paradigmatic example of how a single residue change can alter pharmacology. Mechanistically, S810L increases the stability of the active MR conformation even in the absence of aldosterone. For example, Geller et al. showed that in S810L MR progesterone and 21-hydroxyl-lacking steroids fully activate the receptor. Structural studies have suggested that in S810L MR, the leucine side chain may form lipophilic contacts with H3, stabilizing a conformation that favors H12 closure even in the presence of classical antagonists (Geller et al. 2000; Amazit et al. 2015). It was further proposed that steric crowding between L810 (H5) and Q776 (H3) induces a bending of helix 3 away from helix 5, thereby reshaping the ligand-binding pocket in a more active-like orientation (Pinon et al. 2004).

While it has been proposed that dihydropyridine-type antagonists sterically interfere with H12, the mechanism by which involved helices (H3, H5, and H12) are connected remains elusive (Amazit et al. 2015). Here we show how finerenone modulates the dynamic positioning of H12 compared to eplerenone and how its antagonism persists in the context of disease-associated mutations such as S810L.

## 2 Methods

### 2.1 Model building and Molecular dynamics simulations

The crystal structure of the wild-type human MR-LBD in complex with eplerenone and the coactivator NCOA1 was obtained from the Protein Data Bank (PDB ID: 5MWY) (Bamberg et al. 2018). NCOA1 was removed to ensure unbiased H12 mobility during simulations. We aligned 5MWY with the non-steroidal antagonist-bound MR structure (PDB ID 6L88) (Takahashi et al. 2020). The S810L mutation was introduced using the Protein Builder in MOE (Molecular Operating Environment, 2020.0901 Chemical Computing Group). Finerenone was docked into the LDB of wild-type receptor using GOLD 2023.2.0 (Jones et al. 1997). The docking grid was centered 10 Å around co-crystallized eplerenone. Binding poses were evaluated manually with regard to previously reported interaction profiles (Amazit et al. 2015). Geometry optimizations were performed using MOE to relieve steric clashes between protein side chains. 500 ns MD simulations were performed using Desmond 6.5 with the OPLS-AA force field and explicit TIP3P water model in an orthorhombic box with a 10 Å buffer around the solute protein. Systems were neutralized and supplemented with 0.15 M NaCl. Following default equilibration protocols, production simulations were run in the NPT ensemble at 300 K and 1.01325 bar using the Martyna-Tobias-Klein (MTK) barostat. Three independent replicates were carried out with randomized initial velocities. Trajectory analysis was conducted in VMD version 1.9.4. Distance distributions were visualized as violin plots using the Interactive Dotplot web tool. Median values were determined for each individual replicate and subsequently averaged across triplicates within one system to obtain condition-specific average medians.

### 2.2 Molecular dynamics simulation analysis

Dynamic pharmacophore (dynophore) analysis extracts time-resolved pharmacophoric interaction features from MD trajectories to generate 3D interaction profiles (Bock et al. 2016). The approach has been recently applied to study protein-ligand interactions (Wunsch et al. 2023) We applied dynophore within the LigandScout framework (LigandScout 4.4.3) (Wolber and Langer 2005). In addition to conventional ligand-based dynophores, the method was extended to selected protein side chains (M777 and W806). Therefore, the selected residue was relabeled as “LIG”, allowing LigandScout to recognize it as a pseudo-ligand. We further applied *MDPath* to quantify dynamic residue coupling from the MD trajectories based on normalized mutual information (Doering et al., 2025). We identified and visualized allosteric communication routes within the MR-LBD. For comparison, residues were grouped into AF-2–relevant secondary-structure elements as listed in Table S1.

## 3 Results

### 3.1 Eplerenone stabilizes an agonist-like H12 topology exclusively in S810L MR

In order to elucidate the influence of the S810L mutation on the positioning of H12, we selected W806 (H5) as an AF-2 reference point directly above the binding pocket at the H4-H5 hinge and analyzed the distance between W806 (CD2) and I963 (CB; H12) as a structural indicator for H12 stabilization (Figure 1). While wild-type systems showed median distances of 6.39 Å, the S810L simulations showed divergent values (median distance eplerenone 6.02 Å, median distance finerenone 6.67 Å, Figure 1G-I). Representative frames from the MD simulations suggested a more inward-facing H12 conformation in the eplerenone-S810L complex compared to both, wild-type systems and finerenone-S810L.

**Figure 1.**
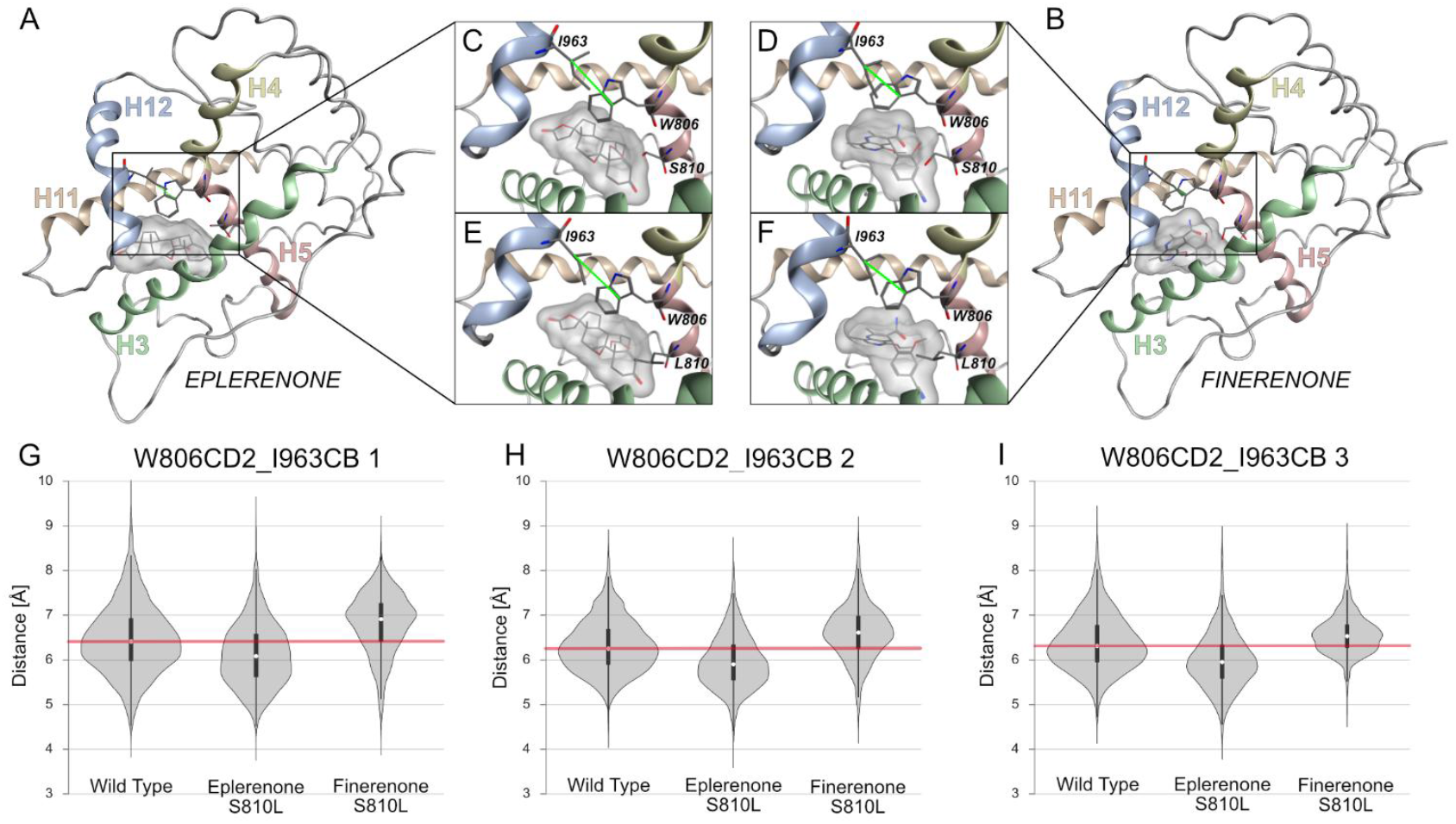
Helix 12 directionality read out by W806–I963 distance across ligands and genotypes. Structural overview of the MR-LBD in complex with eplerenone (A) and finerenone (B). (C-F) Close-up of binding pockets indicating the spatial arrangement of W806, I963 and S/L810 (eplerenone-WT: C, eplerenone-S810L: E; finerenone-WT: D, finerenone-S810L: F). The measured distance between W806CD2 (H5) and I963CB (H12) is indicated in green. (G-I) Violin plots with integrated boxplots of W806CD2–I963CB distance distribution for three independent replicates compare WT with ligand specific S810L-systems. The horizontal red line denotes the combined eplerenone/finerenone wild-type reference-level used for visual comparison. Eplerenone-S810L is shifted consistently towards shorter W806–I963 distances, whereas finerenone-S810L towards longer distances.

### 3.2 Ligand- and mutation-dependent rearrangements of the LBD

We quantified the Cα-Cα distance between M777 (H3) and residue 810 (H5) as surrogate parameter for H3 deformation (Figure 2B). Violin plots summarize the combined distance distribution across all triplicate simulations (Figure 2B). For eplerenone, S810L produces a minor increase in distance. The ligand-dependent difference is already apparent in the wild-type: the steroidal scaffold of eplerenone limits H3-H5 approximation, while finerenone’s planar phenyl ring permits a more compact arrangement. In line with these steric considerations, finerenone shows a smaller M777-810 Cα-Cα distance in wild-type MR compared to the eplerenone complex (7.92 Å vs. 8.39 Å). Moreover, the leucine side-chain of the S810L mutation reorients towards finerenone, further reducing the distance to 7.62 Å (Supplementary Figure S1D).

**Figure 2.**
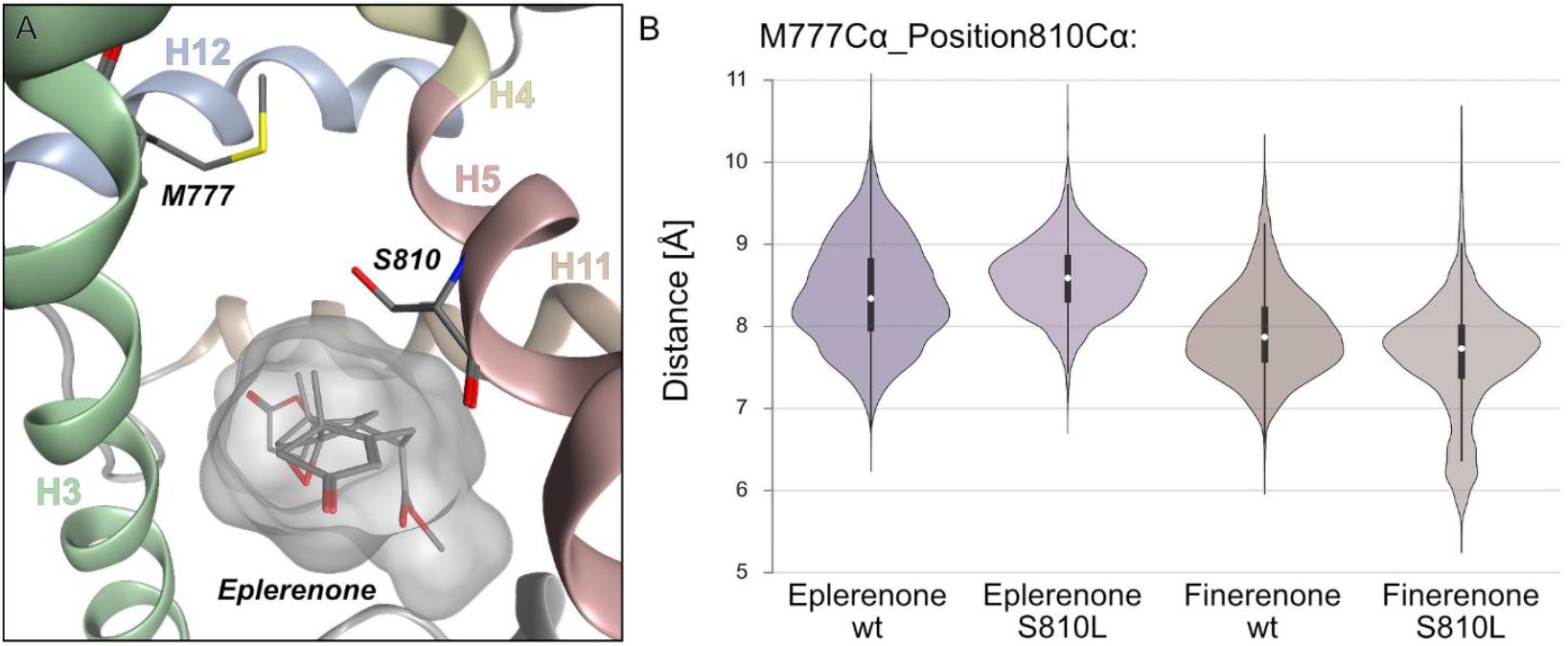
The M777Cα–810Cα distance is used as surrogate for ligand- and mutation-dependent rearrangement at the H3– H5 interface. (A) Close-up of the MR-LBD (PDE:5MWY) bound to eplerenone (grey surface). (B) Violin plots with integrated boxplots of the M777Cα–810Cα distances. Finerenone-WT exhibits shorter distances (7.92± 0.52 Å) than eplerenone-WT (8.39± 0.62 Å) and further compaction for finerenone-S810L (7.62± 0.67 Å), whereas eplerenone-S810L (8.59± 0.43 Å) displays an increase.

### 3.3 Ligand-dependent reorganization of the M777-W806 network in S810L MR

We further analyzed interactions between M777 (H3) and W806 (H5) through residue-centered dynophores in order to collect time-resolved side-chain interaction patterns within AF-2. Reported percentages are averages across triplicates (Figure 3).

**Figure 3.**
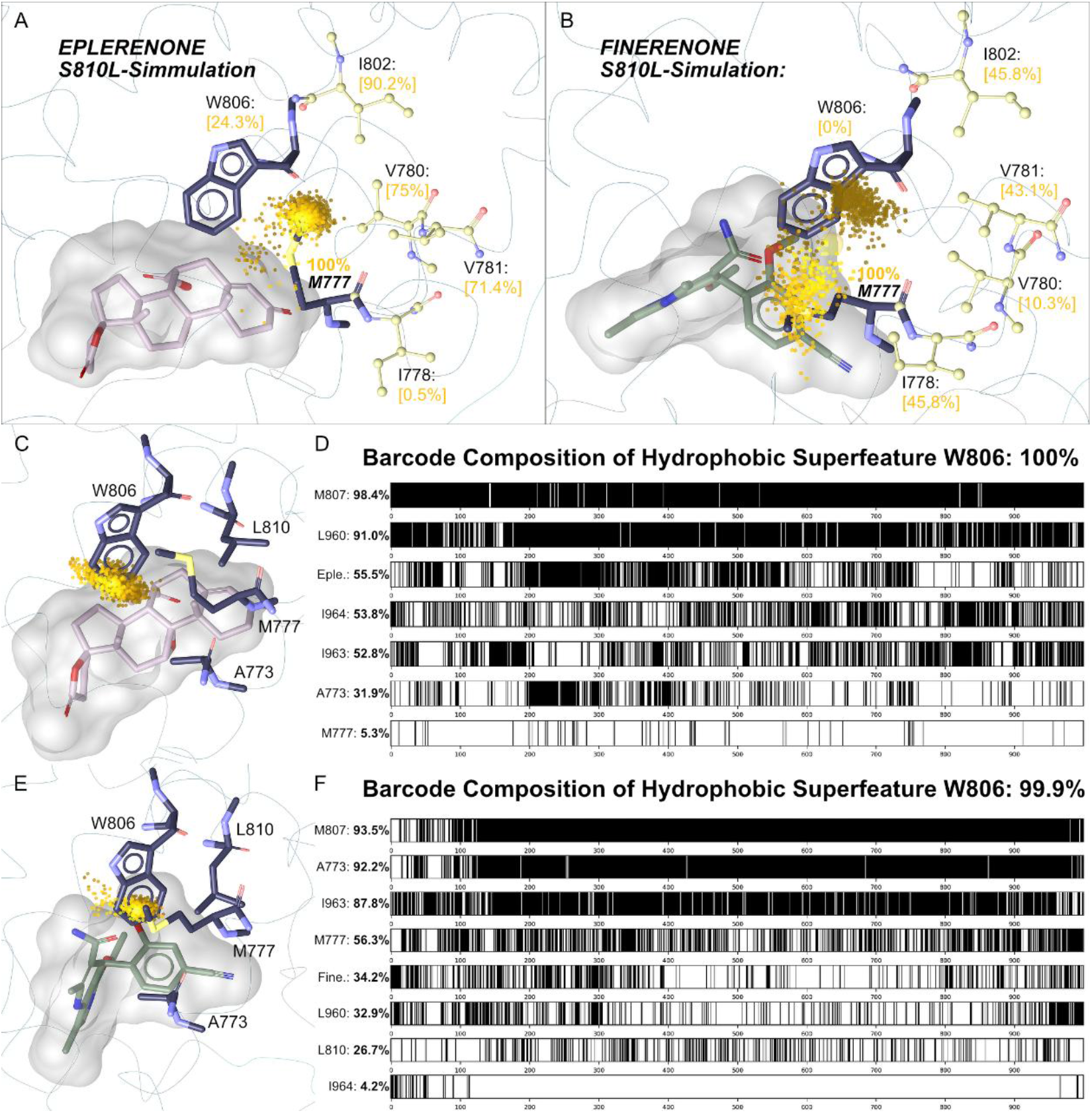
Ligand-dependent reorganization of the M777–W806 network in S810L MR. (A–B) Residue-centered dynophore superfeatures (yellow point clouds) for M777 in eplerenone-S810L (A) and finerenone-S810L (B). AF-2 forming residues on H3/H4 (I802, V780, V781) and W806 H5 as well as I778 H3 are annotated with the fraction of frames in which a hydrophobic contact occurs with M777Cε. In eplerenone-S810L, M777 forms a compact, H3–H4 oriented cloud with high contributions from I802/V780/V781. In finerenone-S810L, density is redistributed towards H3/ligand with increased involvement of I778. (C, E) Residue-centered dynophore superfeatures for W806 in eplerenone-S810L (C) and finerenone-S810L (E). W806 yields a tail-like extension toward H12 residues in eplerenone-S810L, whereas in finerenone-S810L a compact, pocket-oriented cloud directed toward A773 H3. (D, F) Barcode plots of superfeature occurrences for W806-eplerenone-S810L (D) and -finerenone-S810L (F).

In the S810L mutation with bound eplerenone, M777 shows lipophilic contacts with I802 (H4): 83.4%, V781 (H3): 58.5%, V780 (H3): 62.1% and W806 (H5): 24.3% resolved in a sphere-like geometry. V781 is facing away from the binding pocket towards the coactivator binding site and forming a lipophilic pocket together with V780 and I802 (Figure 3A). Within finerenone-S810L systems, this density is redistributed away from the H3-H4 AF-2 (Figure 3B). The superfeature showed lower average occupancies (I802: 58.6%, V781: 50.2%, V780: 35.1% and W806: 13.1%) but additionally showed more frequent contacts with I778 (27% compared to its 4.1% in eplerenone-S810L). V781 is thus able to move towards the original M777 position and occupies the lipophilic pocket, which is formed by I802, V780. M777 also engages H12 residues in both systems, whereas for eplerenone-S810L the contribution is higher (I963: 96.5%, M959: 61.9% and L960: 13.8%) compared to finerenone-S810L (I963: 94%, M959: 36.6% and L960: 0%).

W806, located at the adjacent H5, shifts its lipophilic contacts between H12 and A773 (H3) in a ligand-dependent manner for S810L systems. For eplerenone it maintains strong interactions with H12 residues, yielding 60.9% average occupancy composed of L960 (93.8%), I963 (57.7%) and I964 (31.2%). The W806-A773 contact is less pronounced, averaging 46.6%. In the corresponding dynophore, this pattern is resolved as a tail-like extension directed toward H12 (Figure 3C). In finerenone-S810L, W806-H12 interactions are reduced to an overall occupancy of 47.6% (L960: 53.0%, I963: 76.8%, I964: 13.0%), while the A773 contribution is increased to 65.9% resulting in a more compact superfeature cloud oriented towards the core of the binding pocket and A773 (Figure 3D).

When compared to wild-type systems, the redistribution observed in eplerenone-S810L stands out. For M777, both wild-type systems show either low (eplerenone-WT: 6.8%) or no (finerenone-WT: 0.0%) L960 (H12) interactions. Likewise, W806 maintains a predominant A773 interaction (eplerenone-WT: 75.3%, finerenone-WT: 72.6%), with a level of H12 involvement similar to finerenone-S810L (eplerenone-WT: 41.2%, finerenone-WT: 41.5%) (Supplementary Table S1-S2). Eplerenone-S810L uniquely shifts M777 and W806 towards more robust H12 engagement, while finerenone-S810L keeps wild-type behavior.

### 3.4 Divergent inter-helix interactions are observed for the S810L-eplerenon complex

MDPath-based network analysis revealed distinct communication paths across the investigated ligand– wt/S810L systems (Figure 4 including the number of individual pathways found in the analysis). Communications paths in finerenone WT were predominantly distributed towards helices H3 and H4, with small involvement of H11 and H11–H12 loop. The finerenone S810L system displayed a similar overall distribution, characterized by similar H3 participation and slightly reduced H4 connectivity, while H11 involvement was completely absent and H11-H12 loop directionality was increased. In contrast, eplerenone WT showed weaker H3 and H4 coupling, accompanied by the highest participation of the H3–H4 loop and moderate contributions from the H11–H12 loop. The introduction of the S810L mutation showed a redistribution of communication occurred: path participation moderately increased for H3 and H4, while H3–H4 loop coupling decreased sharply and H11–H12 loop involvement raised substantially. Overall, these results indicate that ligand and genotype jointly shape the preferred allosteric communication routes within the MR-LBD, shifting the balance between H3/H4 connectivity and H11–H12 loop coupling.

**Figure 4.**
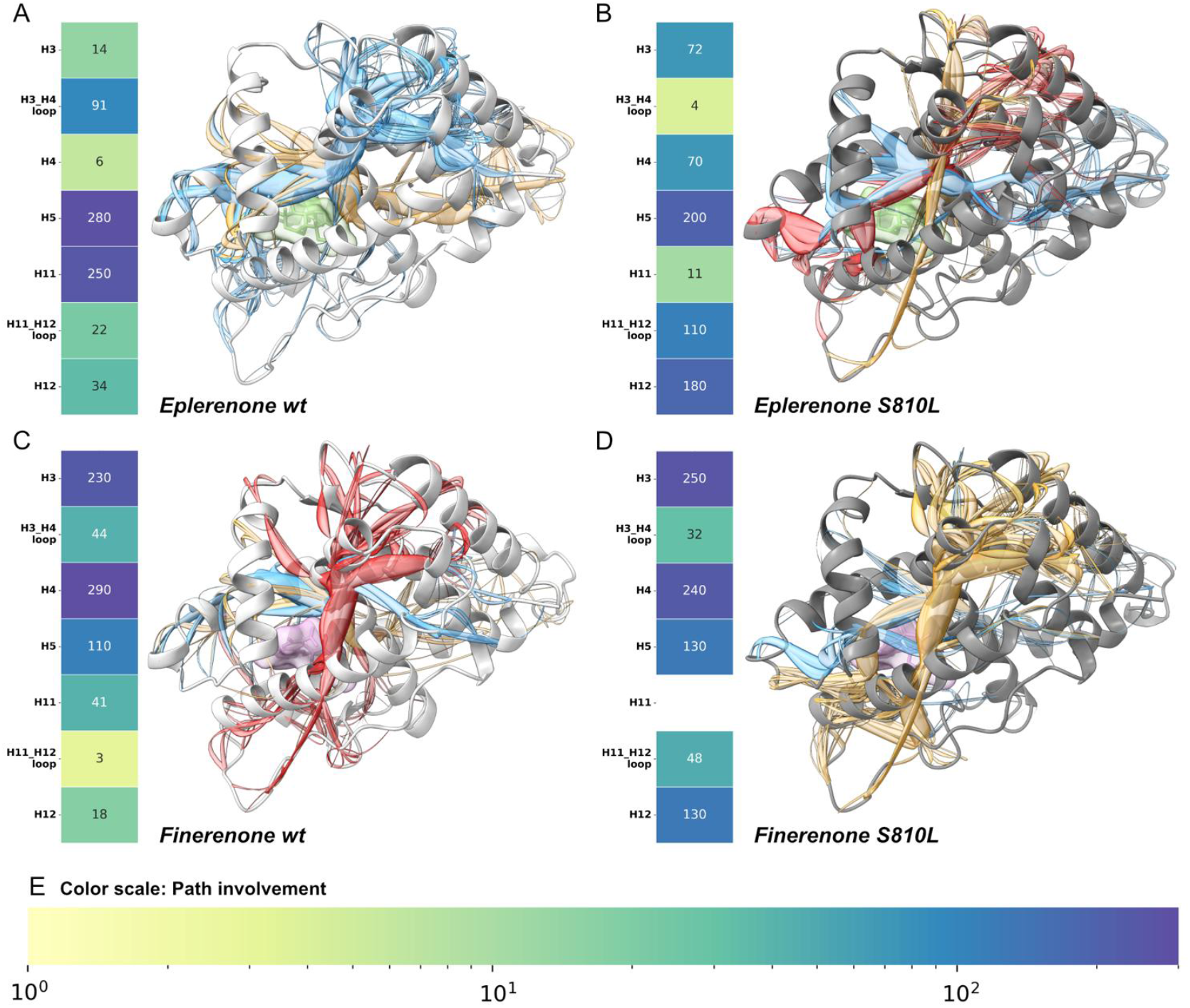
MDPath communication-path mapping in the MR-LBD indicates allosterically linked regions within the MR-LBD across four systems. Corresponding heatmaps indicate the relative participation of residues from AF-2 surface-forming helices H3, H4, H5, H11, H12, and the connecting H3–H4 and H11–H12 loops within the communication networks. (A) Eplerenone-WT, (B) Eplerenone-S810L, (C) Finerenone-WT, and (D) Finerenone-S810L. MDPath-derived communication paths are colored according to their cluster. (C–D) Finerenone systems show enhanced coupling within the AF-2 groove, reflected by high path participation in H3 and H4, accompanied by a reduction in H11 involvement. In contrast, eplerenone systems show a divided pattern: H3 and H4 are moderately engaged, whereas the H11–H12 loop shows enhanced participation in S810L together with a loss of H3–H4 loop involvement. (E) Color scale: Path involvement.

## 4 Discussion

This study addresses how disease-relevant mutations reshape the drug response of the mineralocorticoid receptor. The S810L variant exemplifies this principle by altering AF-2 interactions in a ligand-specific manner: with eplerenone, M777 and W806 reallocate contacts toward H12, whereas with finerenone they pivot toward the binding pocket and A773, maintaining an antagonist-like conformation. Beyond this mutation-centric view, our work characterize finerenone binding in a structurally plausible pose that fits a passive-antagonist template (5MWY) (Bamberg et al. 2018). We reconcile the “bulky/active antagonist” label with prior superpositions showing no H12 clashes by demonstrating that finerenone does not require steric displacement of H12 to destabilize the AF-2 surface (Amazit et al. 2015). In S810L MR, finerenone diverts lipophilic AF-2 side-chain networks away from H12. Accordingly, we extended the classical focus on H3/H5/H12 by incorporating H4-facing contacts and offer residue-level markers for mutation-resilient antagonism. MDPath complements the residue-centered dynophore analysis by showing how ligand-genotype combinations redistribute helix-level communication within AF-2. Across the four systems, finerenone maintains H3/H4-centric routing with low H11/H11-H12 participation. The coupling network remains largely within the binding pocket, rather than H12 hinge-directed and is consistent with the pocket-oriented contact pattern we observed. In contrast, eplerenone S810L exhibits a reallocation away from the H3-H4 loop towards the H11-H12 region, indicating that AF-2 communication converges on the H12 hinge. This routing is compatible with a coactivator-oriented AF-2 arrangement, whereas finerenone retains a pocket-facing communication mode with comparatively weak linkage to H12.

Mechanistically, S810L increases local lipophilicity and reshapes the H5-H3 interface modulating M777 access and orientation. In the eplerenone context, the mutation-induced environment pushes M777 closer to H12 while preserving its ability to engage AF-2-forming residues on H3/H4. The C19 methyl group on eplerenone simultaneously limits deep penetration of W806 toward A773, pushing W806 toward H12 and allowing M777 to bridge the lipophilic H3/H4 cleft to H12. Together, these steric and lipophilic factors consolidate an H12-engaging AF-2 topology in S810L with eplerenone. In contrast, finerenone’s planar phenyl system reorients the L810 side chain toward the ligand compared to eplerenone and compacts the pocket region around M777-810. This eliminates the lipophilic niche seen with eplerenone, moves M777 away from H12, and shifts W806 toward A773, yielding a pocket-oriented topology that withdraws H12 interactions and thereby maintaining antagonism despite the S810L mutation. These contrasts highlight how subtle scaffold differences propagate through S810L-sensitive side-chain networks to produce opposite AF-2 outcomes.

We chose the MR-LBD in complex with eplerenone and the coactivator NCOA1 (PDB ID: 5MWY) as starting point for our modeling, since this structure features a closed H12 in a passive-antagonist conformation (Bamberg et al. 2018). (Supplementary Figure S2).

Our data refine the interpretation of earlier H3 bending proposals for S810L (Pinon et al. 2004). In fully solvated MD settings, we observe a ligand- and mutation-dependent rearrangement at the H3-H5 interface, but these local observations alone do not predict H12 stabilization. This motivated our residue-centered dynophore analysis, which resolves the side-chain traffic that couples the AF-2 constituents and distinguishes H12-engaging.

Two common critiques of MD are worth to mention. First, full H12 transitions are slow, cooperative events that can depend on coactivator pre-organization and are often beyond the sampling window of classical MD. Our simulations were designed to read out residue-network directionality within AF-2 rather than to enforce a full antagonist-like H12 dissociation, which aligns with nuclear-receptor dynamics literature (Batista and Martínez 2013). Second, template choice and starting pose can bias the observations. We addressed this by docking finerenone into a validated MR template (5MWY), documenting backbone compatibility with a non-steroidal complex (6L88), and demonstrating that finerenone produces distinctive, mutation-dependent rearrangements of H3/H4 contacts under identical protocols across independent triplicates. Removing of NCOA1 peptide from 5MWY before MD avoided pre-biasing H12 and ensured comparable, unbiased H12 mobility across ligands and genotypes (Bamberg et al. 2018).

Taken together, these findings place side-chain network reallocations in S810L MR at the center of AF-2 stabilization. Eplerenone uniquely consolidates an H12-engaging topology, whereas finerenone promotes a pocket-oriented topology that helps preserve antagonism in the mutant. Beyond resolving ligand-specific behavior in S810L, this residue-level perspective suggests practical markers for identifying new mutation-resilient antagonists: avoid M777-H12 anchoring and maintain W806-A773 persistence by enabling local narrowing of H3-H5 around M777 and residue 810. Because AF-2 assembly integrates H3, H4, H5 and H12, focusing on network directionality rather than just backbone proximity provides a transferable framework for understanding mutation-altered drug responses across nuclear receptors.

## 6 Author Contributions

All authors have read and approved the final version of the manuscript.

