## Supplemental Information for "Divergent Side-Chain Networks of the Mineralocorticoid Receptor Control Mutation-Altered Drug Response"

**for**

### 1.1 Supplementary Figures

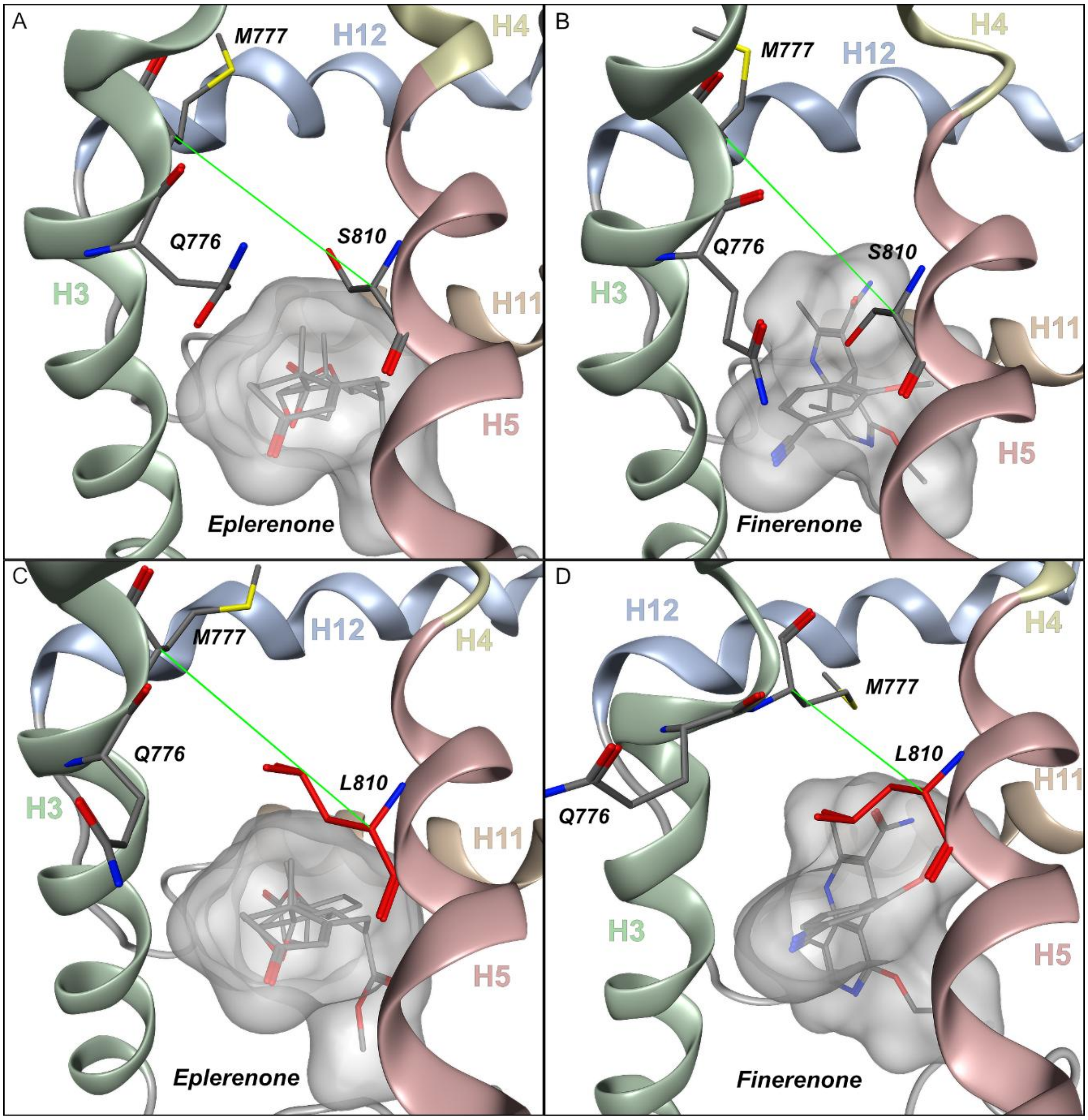


**Supplementary Figure S1.** Representative MD snapshots around the H3–H5–H12 interface.

(A–D) show representative MD simulation frames from the four conditions used in this study: (A) eplerenone–WT, (B) finerenone–WT, (C) eplerenone–S810L, (D) finerenone–S810L. Helices are colored as in the main figures (H3 green, H4 yellow, H5 red, H12 blue; H11 orange). The ligands are shown as sticks with semi-transparent ligand surface. The green vector depicts the M777–(S/L)810 Cα–Cα distance used in Fig. 2B to operationalize H3 bending/approximation. In finerenone-S810L (panel D), L810 adopts a rotamer pointing toward the ligand and compacts the pocket around M777–810, consistent with the reduced Cα–Cα distance reported in Fig. 2B. By contrast, eplerenone–S810L (panel C) maintains a lipophilic niche between H3/H4 and H12, which enables M777 to engage H12 in agreement with Fig. 3.

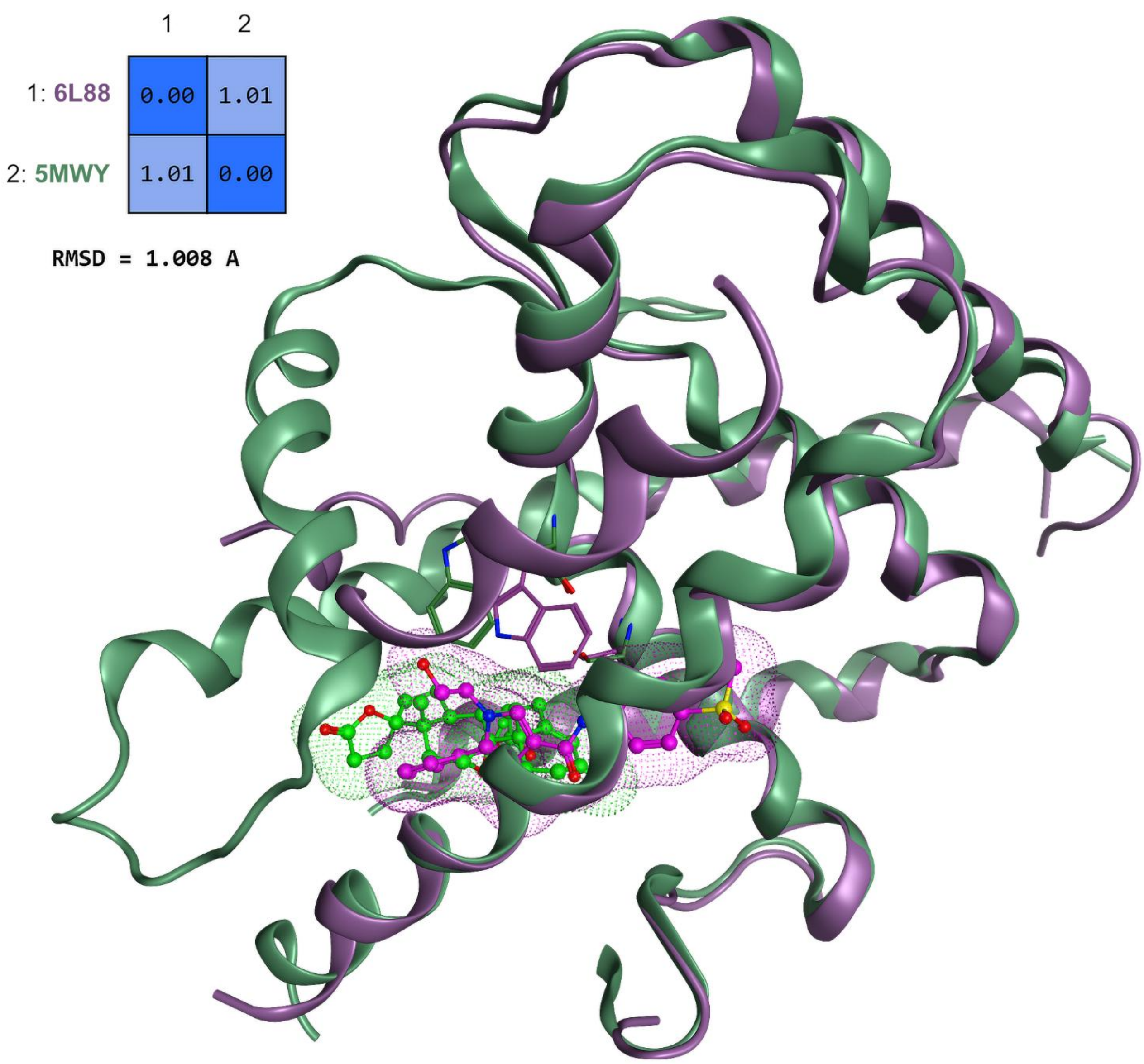


**Supplementary Figure S2.** Structural alignment of MR LBD templates (5MWY vs 6L88).

Superposition of 5MWY (green) and 6L88 (magenta) shows a backbone RMSD ≈ 1.01 Å (matrix inset), indicating a closely matching core fold outside the divergent H11–H12 segment. Esaxerenone (6L88, magenta) occupies a deeper path between H3 and H5 than eplerenone (5MWY, green), correlating with the displaced H12 observed in 6L88. This overlay supports using 5MWY as a neutral starting template for finerenone docking while avoiding esaxerenone-specific AF-2 features (see Methods and Discussion).

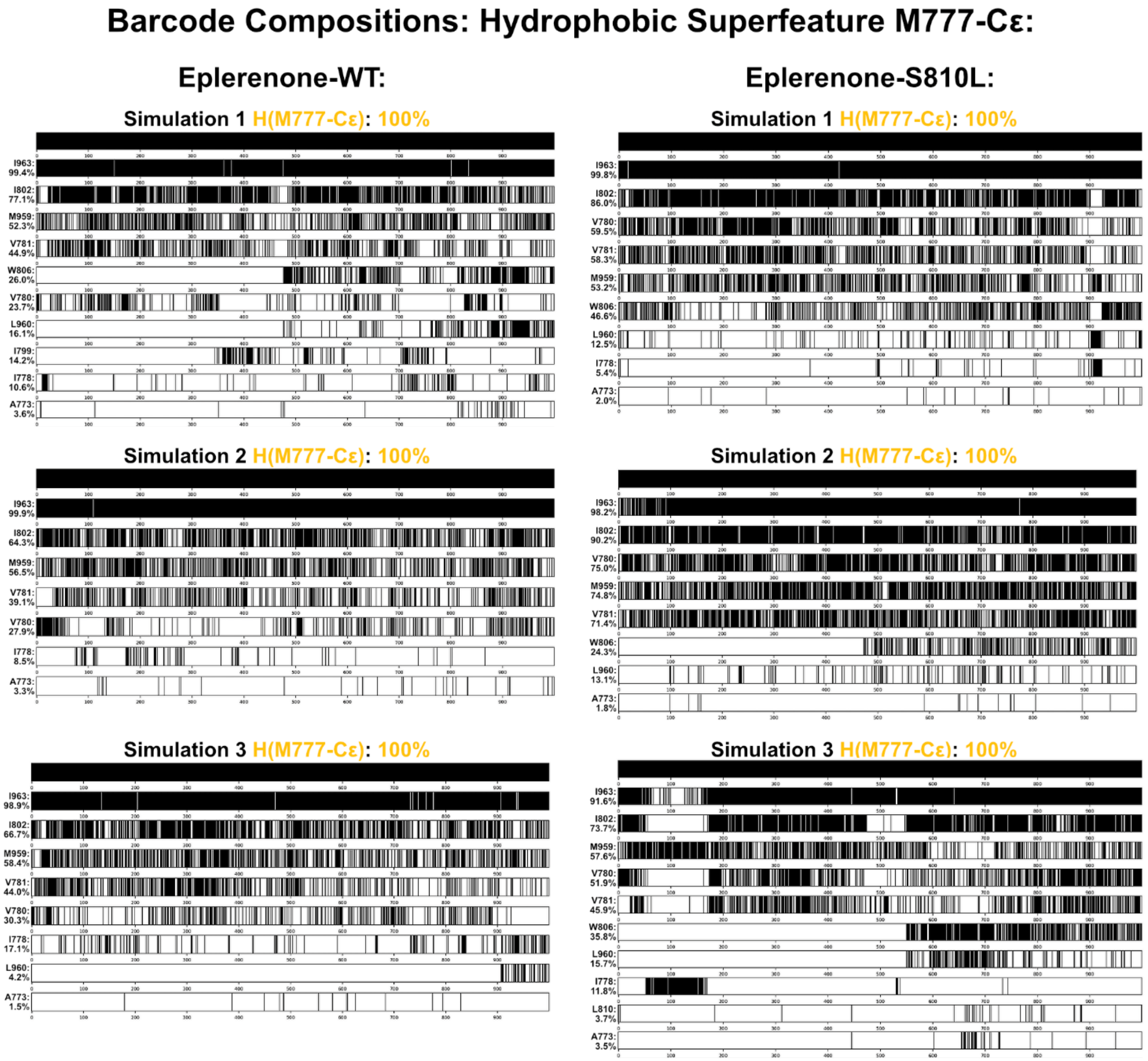


**Supplementary Figure S3.** Eplerenone: M777-Cε hydrophobic contact barcodes.

Hydrophobic barcode compositions of the M777-Cε superfeature for WT (left) and S810L (right), each shown for three independent 500-ns replicates. Rows list interaction partners; each row encodes the trajectory over time, and black segments mark frames in which a hydrophobic contact was detected; percentages give total occupancy per trajectory. Eplerenone–S810L indicates a redistribution toward AF-2-facing partners (I802, V780, V781, W806) and additional H12 engagement (I963, M959, L960) relative to WT, supporting the compact, H12-H3/H4 engaging M777 pattern summarized in Fig. 3A.

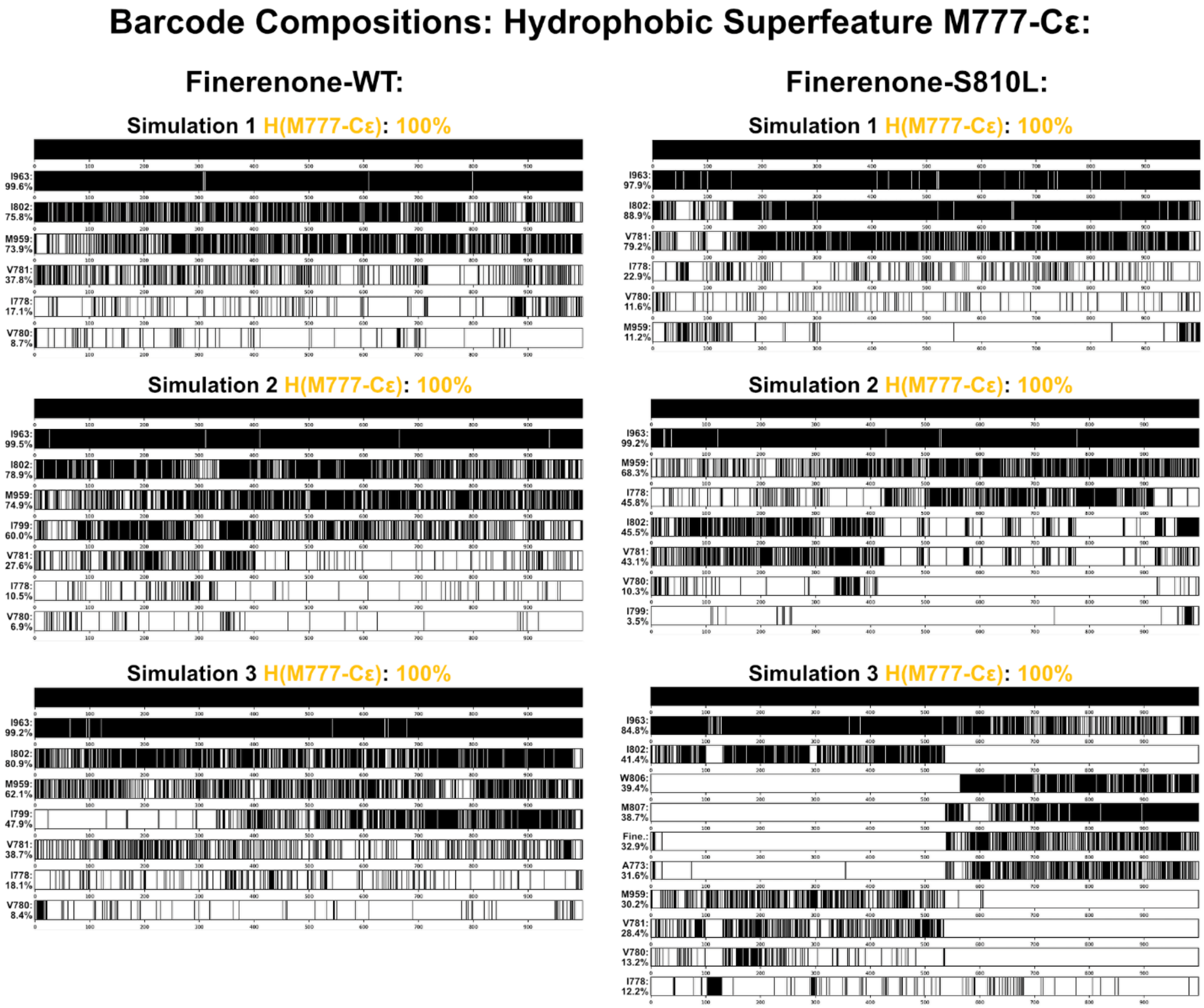


**Supplementary Figure S4.** Finerenone: M777-Cε hydrophobic contact barcodes.

As in S3, for finerenone. Barcodes retain the temporal information as described above. In finerenone–S810L, occupancies at I802, V780, V781, W806 decrease and I778 increases, indicating a pocket-oriented rearrangement. L960 is absent and M959 is reduced, while I963 remains high, matching the M777 patterns in Fig. 3B.

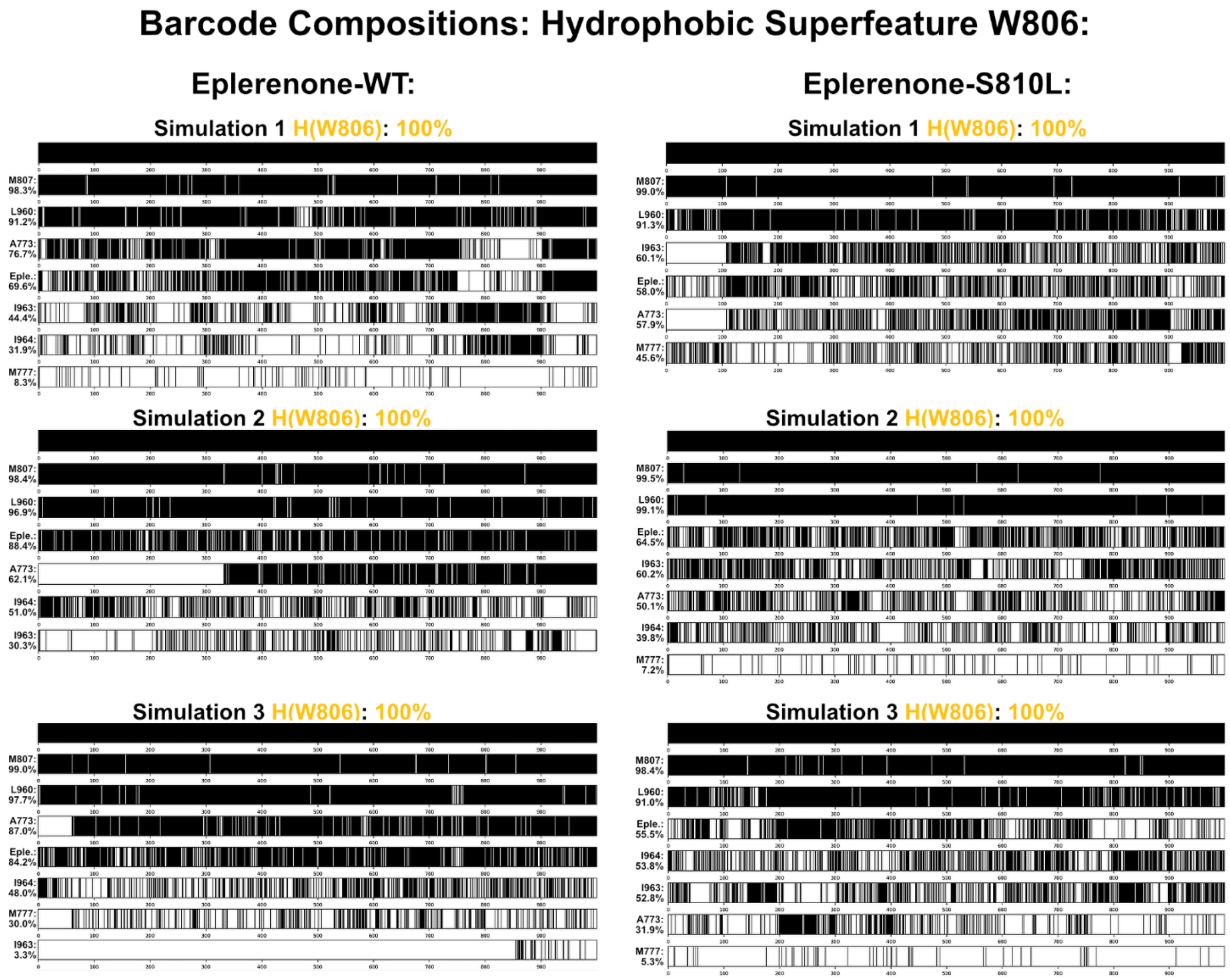


**Supplementary Figure S5.** Eplerenone: W806 hydrophobic contact barcodes.

Hydrophobic barcode compositions of the W806 superfeature for WT (left) and S810L (right), three replicates each. Rows represent the time axis of each trajectory, with black segments indicating frames with hydrophobic contact; percentages report total occupancy. In eplerenone–S810L, H12 contacts (L960, I963, I964) are more persistent and A773 is reduced relative to WT, reflecting a routing of W806 toward H12 that complements the M777 findings and supporting H12 engaging W806 pattern (Fig. 3C). WT retains predominant A773 engagement with comparatively weaker H12 involvement.

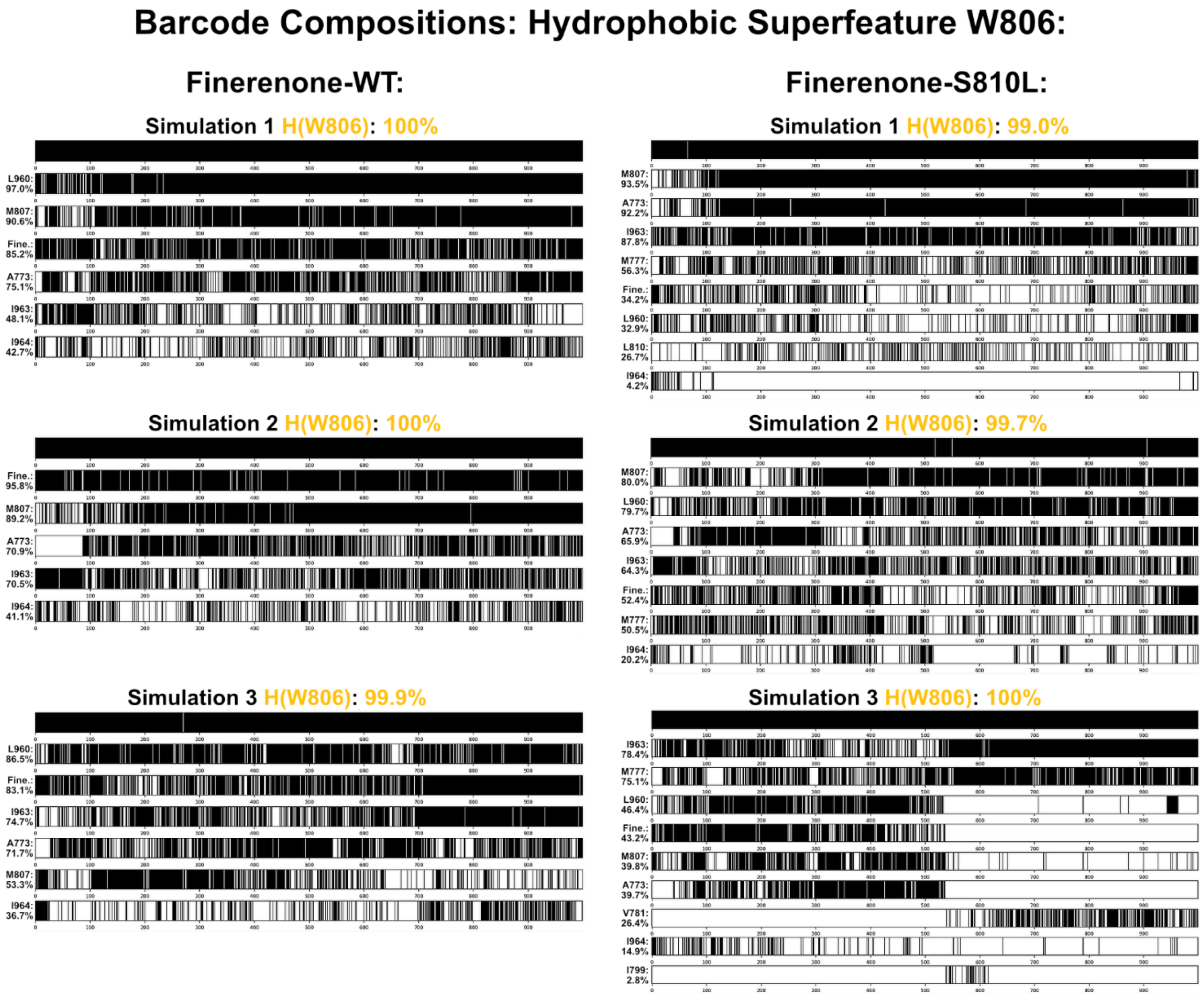


**Supplementary Figure S6.** Finerenone: W806 hydrophobic contact barcodes.

As in S5, for finerenone, with the same time-resolved barcode convention. In finerenone–S810L, A773 becomes the dominant partner and H12 contacts are overall weaker, consistent with a pocket-oriented topology. The WT pattern resembles finerenone–S810L rather than eplerenone–S810L (Fig. 3D).

***Supplementary Table S1. M777-Cε hydrophobic superfeature: partner occupancies (%) across triplicate simulations for eplerenone and finerenone in WT and S810L MR.***

| **Eplerenone-WT**: M777 Hydrophobic Superfeature Occupancy | | | | | | |
|---|---|---|---|---|---|---|
| | | | Simulation | | | |
| Helix | Residue | 1 | 2 | 3 | Averaged | |
| 5 | W806 | 26.0% | 0% | 0% | 8.7% | |
| 4 | I802 | 77.1% | 64.3% | 66.7% | 69.4% | |
| | V781 | 44.9% | 39.1% | 44.0% | 42.7% | |
| 3 | V780 | 23.7% | 27.9% | 30.3% | 27.3% | 27.4% |
| | I778 | 10.6% | 8.5% | 17.1% | 12.1% | |
| | I963 | 99.4% | 99.9% | 98.9% | 99.4% | |
| 12 | L960 | 16.1% | 0% | 4.2% | 6.8% | 54.0% |
| | M959 | 52.3% | 56.5% | 58.4% | 55.7% | |
| **Eplerenone-S810L**: M777 Hydrophobic Superfeature Occupancy | | | | | | |
| | | | Simulation | | | |
| Helix | Residue | 1 | 2 | 3 | Averaged | |
| 5 | W806 | 46.4% | 24.3% | 35.8% | 35.5% | |
| 4 | I802 | 86.0% | 90.2% | 73.7% | 83.3% | |
| | V781 | 58.3% | 71.4% | 45.9% | 58.5% | |
| 3 | V780 | 59.5% | 75.0% | 51.9% | 62.1% | 43.3% |
| | I778 | 5.4% | 0% | 11.8% | 9.3% | |
| | I963 | 99.8% | 98.2% | 91.6% | 96.5% | |
| 12 | L960 | 12.5% | 13.1% | 15.7% | 13.8% | 57.4% |
| | M959 | 53.2% | 74.8% | 57.6% | 61.9% | |
| **Finerenone-WT**: M777 Hydrophobic Superfeature Occupancy | | | | | | |
| | | | Simulation | | | |
| Helix | Residue | 1 | 2 | 3 | Averaged | |
| 5 | W806 | 0% | 0% | 0% | 0% | |
| 4 | I802 | 75.8% | 78.9% | 80.9% | 78.5% | |
| | V781 | 37.8% | 27.6% | 38.7% | 34.7% | |
| 3 | V780 | 8.7% | 6.9% | 8.4% | 8% | 19.3% |
| | I778 | 17.1% | 10.5% | 18.1% | 15.2% | |
| | I963 | 99.6% | 99.5% | 99.2% | 99.4% | |
| 12 | L960 | 0% | 0% | 0% | 0% | 56.6% |
| | M959 | 73.9% | 74.9% | 62.1% | 70.3% | |
| **Finerenone-S810L**: M777 Hydrophobic Superfeature Occupancy | | | | | | |
| | | | Simulation | | | |
| Helix | Residue | 1 | 2 | 3 | Averaged | |
| 5 | W806 | 0% | 0% | 39.4% | 13.1% | |
| 4 | I802 | 88.9% | 45.5% | 41.4% | 58.6% | |
| | V781 | 79.2% | 43.1% | 28.4% | 50.2% | |
| 3 | V780 | 11.6% | 10.3% | 13.2% | 11.7% | 29.6% |
| | I778 | 22.9% | 45.8% | 12.2% | 27.0% | |
| | I963 | 97.9% | 99.2% | 84.8% | 94.0% | |
| 12 | L960 | 0% | 0% | 0% | 0% | 43.5% |
| | M959 | 11.2% | 68.3% | 30.2% | 36.6% | |

Each block lists WT and S810L systems; columns (1–3) are individual 500-ns replicates, (Averaged) is the mean occupancy across replicates. Rows denote interaction partners grouped by helix (H5: W806; H4: I802; H3: V781, V780, I778; H12: I963, L960, M959). An entry reflects the fraction of frames with a hydrophobic contact detected by residue-centered dynophore for M777. The tabulated percentages provide a compact overview of raw values to facilitate cross-system comparison and correspond to the percentages summarized in the main text and support Fig. 3A–B.

***Supplementary Table S2. W806 hydrophobic superfeature: partner occupancies (%) across triplicate simulations for eplerenone and finerenone in WT and S810L MR.***

| **Eplerenone-WT**: W806 Hydrophobic Superfeature Occupancy | | | | | | |
|---|---|---|---|---|---|---|
| | | | Simulation | | | |
| Helix | Residue | 1 | 2 | 3 | Averaged | |
| 3 | A773 | 76.7% | 62.1% | 87.0% | 72.3% | |
| | I964 | 31.9% | 51.0% | 48.0% | 43.6% | |
| 12 | I963 | 44.4% | 30.3% | 3.3% | 26.0% | 55.0% |
| | L960 | 91.2% | 96.9% | 97.7% | 95.3% | |
| **Eplerenone-S810L**: W806 Hydrophobic Superfeature Occupancy | | | | | | |
| | | | Simulation | | | |
| Helix | Residue | 1 | 2 | 3 | Averaged | |
| 3 | A773 | 57.9% | 50.1% | 31.9% | 46.6% | |
| | I964 | 0% | 39.8% | 53.8% | 31.2% | |
| 12 | I963 | 60.1% | 60.2% | 52.8% | 57.7% | 60.9% |
| | L960 | 91.3% | 99.1% | 91% | 93.8% | |
| **Finerenone-WT**: W806 Hydrophobic Superfeature Occupancy | | | | | | |
| | | | Simulation | | | |
| Helix | Residue | 1 | 2 | 3 | Averaged | |
| 3 | A773 | 75.1% | 70.9% | 71.1% | 72.4% | |
| | I964 | 42.7% | 41.1% | 36.7% | 40.2% | |
| 12 | I963 | 48.1% | 70.5% | 74.7% | 64.4% | 55.2% |
| | L960 | 97% | 0% | 86.5% | 61.2% | |
| **Finerenone-S810L**: W806 Hydrophobic Superfeature Occupancy | | | | | | |
| | | | Simulation | | | |
| Helix | Residue | 1 | 2 | 3 | Averaged | |
| 3 | A773 | 92.2% | 65.9% | 39.7% | 65.9% | |
| | I964 | 4.2% | 20.2% | 14.9% | 13.1% | |
| 12 | I963 | 87.8% | 64.3% | 78.4% | 76.8% | 47.6% |
| | L960 | 32.9% | 79.7% | 46.4% | 53.0% | |

Blocks and columns as in S1. Rows list interaction partners A773 (H3) and H12 residues (I964, I963, L960). Occupancies report the fraction of frames with a hydrophobic contact detected by residue-centered dynophore for W806. These raw percentages provide a concise cross-condition summary and underpin the W806 patterns discussed in the main text and Fig. 3C–D (H12-engaging vs pocket-oriented routing).

**Supplementary Table S3. Residue mapping used for AF-2–related secondary-structure grouping in MDPath analyses**

| Secondary-structure element | Residues included |
|---|---|
| H3 | V781, M777, A773, N770 |
| H3-H4 loop | F790 |
| H4 | I802 |
| H5 | S/L810, W806, R817 |
| H11 | T945, C942 |
| H11–H12 loop | F956, E955, V954 |
| H12 | Q967, I964, I963, L960 |
